# Over-expression of the RieskeFeS protein increases electron transport rates and yield in *Arabidopsis*

**DOI:** 10.1101/133702

**Authors:** Andrew J. Simkin, Lorna McAusland, Tracy Lawson, Christine A. Raines

## Abstract

In this study we have generated transgenic Arabidopsis plants over-expressing the Rieske FeS protein (PetC), a component of the cytochrome *b*_*6*_*f* (cyt *b*_*6*_*f*) complex. Increasing the levels of this protein, resulted in the concomitant increase in the levels of cyt *f* (PetA) and cyt *b*_*6*_ (PetB), core proteins of the cyt *b*_*6*_*f* complex. Interestingly, an increase in the levels of proteins in both the PSI and PSII complexes was also seen in the Rieske FeS ox plants. Although the mechanisms leading to these changes remain to be identified, the transgenic plants presented here provide novel tools to explore this. Importantly, the overexpression of the Rieske FeS protein resulted in a substantial and significant impact on the quantum efficiency of PSI and PSII, electron transport, biomass and seed yield in Arabidopsis plants. These results demonstrate the potential for manipulating electron transport processes to increase crop productivity.

**One-sentence summary:** Over-expression of the Rieske FeS protein results in significant increases in the quantum efficiencies or PSI and PSII, increases in A_max_ and has the potential to increase crop productivity

## Introduction

Increasing food and fuel demands by the growing world population has led to the need to develop higher yielding crop varieties (Fischer and Edmeades, 2010; Ray et al., 2012). Transgenic studies, modelling approaches and theoretical considerations provide evidence that increasing photosynthetic capacity is a viable route to increase the yield of crop plants (Zhu et al., 2010; Raines, 2011; Long et al., 2015; von Caemmerer and Furbank, 2016). There is now a growing body of experimental evidence showing that increasing the levels of photosynthetic enzymes in carbon metabolism, results in increased photosynthesis and plant biomass (Miyagawa et al., 2001; Raines, 2006, 2011; Lefebvre et al., 2005; Rosenthal et al., 2011; Uematsu et al., 2012; Simkin et al., 2015, 2017; Driever et al., 2017). In addition, manipulation of photosynthetic electron transport by introduction of the algal cytochrome c6 protein has been shown to improve the efficiency of photosynthesis and to stimulate plant growth in low light (Chida et al., 2007). One endogenous target identified for manipulation is the cytochrome *b*_*6*_*f* (cyt *b*_*6*_*f*) complex which is located in the thylakoid membrane and functions in both linear and cyclic electron transport, providing ATP and NADPH for photosynthetic carbon fixation. Initially, cyt *b*_*6*_*f* inhibitors (Kirchhoff et al., 2000) and later transgenic antisense studies suppressing the accumulation of the Rieske FeS protein (PetC), a component of the cyt *b*_*6*_*f* complex, have demonstrated that the activity of the cyt *b*_*6*_*f* complex is a key determinant of the rate of electron transport (Price et al., 1995, 1998; Anderson et al., 1997; Ruuska et al., 2000; Yamori et al., 2011a,b).

The finding that the cyt *b*_*6*_*f* complex is a potential limiting step in the electron transport chain suggests that by increasing the activity of this complex it may be possible to increase the rate of photosynthesis. However, questions have been raised about the feasibility of manipulating this multiprotein, membrane located, complex given that it is composed of eight different subunits, six being encoded in the chloroplast genome (PetA (cyt *f*), PetB (cyt *b*_*6*_), PetD, PetG, PetL and PetN) and two in the nucleus (PetC, (Rieske FeS) and PetM) (Willey and Grey, 1988; Anderson et al., 1992; Knight et al., 2002; Cramer & Zhang, 2006, Cramer et al., 2006; Baniulis et al., 2009; Schöttler et al., 2015). Furthermore, this protein complex functions as a dimer with the transmembrane domains of both the Rieske FeS and cyt *b*_6_ proteins being directly implicated in the monomer–monomer interaction and stability of the complex and the *petD* gene product functioning as a scaffold (Hager et al., 1999; Schwenkert et al., 2007; Hojka et al., 2014; Cramer et al., 2006). Essential roles in the assembly and stability of the cyt *b*_*6*_*f* complex have also been shown for the PetG, PetN and PetM subunits and a minor role in stability assigned to the PetL gene product (Schöttler et al., 2007; Bruce and Malkin, 1991; Kuras and Wollman, 1994; Hager et al., 1999; Monde et al., 2000; Schwenkert et al., 2007; Hojka et al., 2014).

Notwithstanding both the genetic and structural complexity of the cyt *b*_*6*_*f* complex, it has been shown previously that it is possible to manipulate the levels of the cyt *b*_*6*_*f* complex by down regulation of the expression of the Rieske FeS protein (Price. et al., 1998; Yamori et al., 2011a). It has also been shown that the Rieske FeS protein is one of the subunits required for the successful assembly of the cyt *b*_*6*_*f* complex (Miles, 1982; Metz et al., 1983; Barkan et al., 1986; Anderson et al., 1997). Based on these results, we reasoned that over-expression of the Rieske FeS protein could be a feasible approach to take in order to increase the electron flow through the cyt *b*_*6*_*f*. In this paper we report on the production of Arabidopsis with increased levels of the tobacco Rieske FeS protein and we show that this manipulation resulted in an increase in photosynthetic electron transport, CO_2_ assimilation and yield. This work provides evidence that the process of electron transport is potential route for the improvement of plant productivity.

## Material and methods

### Rieske iron sulphur protein of the cytochrome *b*_*6*_*f* (*cyt b*_*6*_*f*)

The full-length coding sequence of the Rieske iron sulphur protein of the cytochrome *b*_*6*_*f (*X64353) was amplified by RT-PCR using primers NtRieskeFeSF (5’cacc<u>ATGG</u>CTTCTTCTACTCTTTCTCCAG’3) and NtRieskeFeSR (5’<u>CTA</u>AGCCCACCATGGATCTTCACC’3). The resulting amplified product was cloned into pENTR/D (Invitrogen, UK) to make pENTR-*NtRieskeFeS* and the sequence was verified and found to be identical. The full-length cDNA was introduced into the pGWB2 gateway vector (Nakagawa et al., 2007: AB289765) by recombination from the pENTR/D vector to make pGW-NtRieske (B2-NtRi). cDNA are under transcriptional control of the 35s tobacco mosaic virus promoter, which directs constitutive high-level transcription of the transgene, and followed by the *nos* 3' terminator. Full details of the B2-NtRi construct assembly can be seen in Supplemental Fig. S1.

### Generation of transgenic plants

The recombinant plasmid B2-NtRi was introduced into wild type Arabidopsis by floral dipping (Clough and Bent, 1998) using *Agrobacterium tumefaciens* GV3101. Positive transformants were regenerated on MS medium containing kanamycin (50mg L^-1^), hygromycin (20mg L^-1^). Kanamycin/hygromycin resistant primary transformants (T1 generation) with established root systems were transferred to soil and allowed to self-fertilize.

### Plant Growth Conditions

Wild type T2 Arabidopsis plants resulting from self-fertilization of transgenic plants were germinated in sterile agar medium containing Murashige and Skoog salts, selected on kanamycin and grown to seed in soil (Levington F2, Fisons, Ipswich, UK) and lines of interest were identified by western blot and qPCR. For experimental study, T3 progeny seeds from selected lines were germinated on soil in controlled environment chambers at an irradiance of 130 μmol m^-2^ s^-1^ in an 8 h/16 h square-wave photoperiod, with an air temperature of 22°C and a relative humidity of 60%. Plants position was randomised and the position of the trays rotated daily under the light. Leaf areas were calculated from photographic images using ImageJ software (imagej.nih.gov/ij). Wild type plants used in this study were a combined group of WT and null segregants from the transgenic lines, verified by PCR for non-integration of the transgene, as no significant differences in growth parameters were seen between them (Supplemental Fig. S2).

### Protein Extraction and Immunoblotting

Four leaf discs (0.6-cm diameter) from two individual leaves were taken, immediately plunged into liquid N^2^ and subsequently stored at −80°C. Samples were ground in liquid nitrogen and protein quantification determined (Harrison et al., 1998). Samples were loaded on an equal protein basis, separated using 12% (w/v) SDS-PAGE, transferred to polyvinylidene difluoride membrane, and probed using antibodies raised against the cytochrome *b*_*6*_*f complex* proteins cyt *f* (PetA: (AS08306), cyt *b*_*6*_ (PetB: (AS03034), Rieske FeS (PetC: AS08330), the photosystem I Lhca1 (AS01005) and PsaA (AS06172) proteins the Photosystem II PsbA/D1 (AS01016) and PsbD/D2 (AS06146) proteins, ATP synthase delta subunit (AS101591), and against the glycine decarboxylase H-subunit (AS05074), all purchased from Agrisera (via Newmarket Scientific UK). FBPA antibodies were raised against a peptide from a conserved region of the protein [C]-ASIGLENTEANRQAYR-amide, Cambridge Research Biochemicals, Cleveland, UK (Simkin et al., 2015). Proteins were detected using horseradish peroxidase conjugated to the secondary antibody and ECL chemiluminescence detection reagent (Amersham, Buckinghamshire, UK). Proteins were quantified using a Fusion FX Vilber Lourmat imager (Peqlab, Lutterworth, UK) as previously described (Vialet-Chabrand et al., 2017).

### Chlorophyll Fluorescence Imaging

Chlorophyll fluorescence measurements were performed on 10-day-old Arabidopsis seedlings that had been grown in a controlled environment chamber at a photosynthetic photon flux density (PPFD) of 130 μmol m^-2^s^-1^ ambient CO_2_ at 22°C. Images of the operating efficiency of photosystem two (PSII) photochemistry, (*F*_q_’/*F*_m_’) were taken at PPFDs of 310 and 600 μmol m^-2^ s^-1^ using a chlorophyll fluorescence imaging system, (Technologica, Colchester, UK; Barbagallo et al., 2003; Baker and Rosenqvist, 2004). *F*_q_’/*F*_m_’, was calculated from measurements of steady state fluorescence in the light (*F*’) and maximum fluorescence in the light (*F*_m_’) was obtained after a saturating 800 ms pulse of 6200 μmol m^-2^ s^-1^ PPFD using the following equation *F*_q_’/*F*_m_’ = (*F*_m_’-*F*’)/*F*_m_’. (Baker et al., 2001; Oxborough and Baker 1997a).

### A/C_i_ response curves

The response of net photosynthesis (*A*) to intracellular CO_2_ (*C*_i_) was measured using a portable gas exchange system (CIRAS-1, PP Systems Ltd, Ayrshire, UK). Leaves were illuminated using a red-blue light source attached to the gas-exchange system, and light levels were maintained at saturating photosynthetic photon flux density (PPFD) of 1000 μmol m^-2^ s^-^ 1 with an integral LED light source (PP Systems Ltd, Ayrshire, UK) for the duration of the *A*/*C*_i_ response curve. Measurements of *A* were made at ambient CO_2_ concentration (*C*_a_) of 400 μmol mol^-1^, before *C*_a_ was decreased in a stepwise manner to 300, 200, 150, 100, 50 μmol mol^-1^ before returning to the initial value and increased to 500, 600, 700, 800, 900, 1000, 1100, 1200 μmol mol^-1^. Leaf temperature and vapour pressure deficit (VPD) were maintained at 22°C and 1 ± 0.2 kPa respectively. The maximum rates of Rubisco-(*Vc*_*max*_) and the maximum rate of electron transport for RuBP regeneration (*J*_*max*_) were determined and standardized to a leaf temperature of 25°C based on equations from Bernacchi et al. (2001), and McMurtrie and Wang (1993) respectively.

### Photosynthetic capacity

Photosynthesis as a function of PPFD (*A*/*Q* response curves) was measured using a Li-Cor 6400XT portable gas exchange system (Li-Cor, Lincoln, Nebraska, USA). Cuvette conditions were maintained at a leaf temperature of 22°C, relative humidity of 50-60%, and ambient growth CO_2_ concentration 400 μmol mol^-1^ for plants grown in ambient conditions). Leaves were initially stabilized at saturating irradiance 1000 μmol m^-2^ s^-1^, after which *A* and *g*_s_ was measured at the following PPFD levels; 0, 50, 100, 150, 200, 250, 300, 350, 400, 500, 600, 800, 1000 μmol m^-2^ s^-1^. Measurements were recorded after *A* reached a new steady state (1-3 min) and before stomatal conductance (*g*_s_) changed to the new light levels. *A*/*Q* analyses were performed at 21% and 2% [O_2_].

### PSI and PSII quantum efficiency

The photochemical quantum efficiency of PSII and PSI in transgenic and WT plants was measured following a dark-light induction transition using a Dual-PAM-100 instrument (Walz, Effeltrich, Germany) with a DUAL-DR measuring head. Plants were dark adapted for 20 min before placing in the instrument. Following a dark adapted measurement plants were illuminated with 220 μmol m^-2^ s^-1^ PPFD. The maximum quantum yield of PSII was measured following a saturating pulse of light for 600 ms saturating pulse of light at an intensity of 6200 μmol m^-2^ s^-1^. The PSII operating efficiency was determined as described by the routines above. PSI quantum efficiency was measured as an absorption change of P700 before and after a saturating pulse of 6200 μmol m^-2^ s^-1^ for 300 ms (which fully oxidizes P700) in the presence of far-red light with a FR pre-illumination of 10s. Both measurements were recorded every minute for 5 min). *q*_p_ or (*F*_v_’/*F*_m_’), was calculated from measurements of steady state fluorescence in the light (*F*’) and maximum fluorescence in the light (*F*_m_’) whilst minimal fluorescence in the light (*F*_*o*_’) was calculated following the equation of Oxborough and Baker (1997b). The fraction of open PSII centres (*q*_L_) was calculated from *q*_p_ x *F*_*o*_*’*/*F* (Baker, 2008).

### Pigment extraction and HPLC analysis

Chlorophylls and carotenoids were extracted using n,n-dimethylformamide (DMF) as previously described (Inskeep and Bloom, 1985), which was subsequently shown to suppressed chlorophyllide formation in Arabidopsis leaves (Hu et al., 2013). Briefly, two leaf discs collected from two different leaves were immersed in DMF at 4°C for 48 hours and separated by UPLC as described by Zapata et al. (2000).

### Leaf Thickness

Leaves of equivalent developmental stage were collected from plants after 28 days of growth. Strips were cut from the centre of the leaf, avoiding the mid-vein, preserved in 5% glutaraldehyde, stored at 4°C for 24 h followed by dehydration in sequential ethanol solutions of 20, 40, 80 and 100%. The samples were placed in LR white acrylic resin (Sigma-Aldrich, Gillingham, UK), refrigerated for 24 h, embedded in capsules and placed at 60°C for 24 h. Sections (0.5μm) were cut using a Reichert-Jung Ultracut microtome (Ametek Gmbh, Munich, Germany), fixed, stained and viewed under a light microscope (Lopez-Juez et al., 1998). Leaf thickness was determined by measuring leaves from two to three plants from line 9 and 11 compared to leaves from four wild type plants.

### Statistical Analysis

All statistical analyses were done by comparing ANOVA, using Sys-stat, University of Essex, UK. The differences between means were tested using the Post hoc Tukey test (SPSS, Chicago).

## Results

### Production and selection of Rieske FeS ox transformants

The full-length tobacco Rieske FeS coding sequence from the cyt *b*_*6*_*f* complex was used to generate an over-expression construct B2-NtRi (Supplemental Fig. S1). Following floral dipping, transgenic Arabidopsis plants were selected on both kanamycin/hygromycin containing medium (Nakagawa et al., 2007) and plants expressing the integrated transgenes identified using RT-PCR (data not shown). Proteins were extracted from leaves of the T1 progeny allowing the identification of three lines with increased levels of the Rieske FeS protein (PetC) (Supplemental Fig. S3A). Immunoblot analysis of T3 progeny of lines 9, 10 and 11 were shown to have higher levels of the Rieske FeS protein when compared to wild type (WT) (Fig. 1 and Supplemental Fig. S3B). The over-expression of the Rieske FeS protein (hereafter referred to as Rieske FeS ox) resulted in a concomitant increase in both cyt *f* (PetA) and cyt *b*_*6*_ (PetB) (Fig. 1A). An increase in the level of the PSI type I chlorophyll a/b-binding protein (LhcaI) and an increase in the core protein of PSI (PsaA) was also observed. Furthermore, the D1 (PsbA) and D2 (PsbD) proteins which form the reaction centre of PSII were also shown to be elevated in Rieske FeS ox lines. Finally, an increase in the ATP synthase delta subunit (AtpD) was also observed in Rieske FeS ox lines (Fig. 1A). In contrast, no notable differences in protein levels for either the chloroplastic FBP aldolase (FBPA), the mitochondrial glycine decarboxylase-H protein (GDC-H) or the Rubisco large subunit were observed (Fig. 1A). A quantitative estimate of the changes in protein levels was determined from the immunoblots of leaf extracts isolated from two to three independent plants per lines. An example is shown in Fig. 1B. These results showed a 2-2.5 fold increase in the Rieske Fe S protein relative to WT plants and a similar increase was also observed for cyt *f*, cyt *b*_*6*_, Lhca1, D2 and PsaA (Fig 1C). No increase in the stromal FBPA protein was evident.

**Figure 1.**
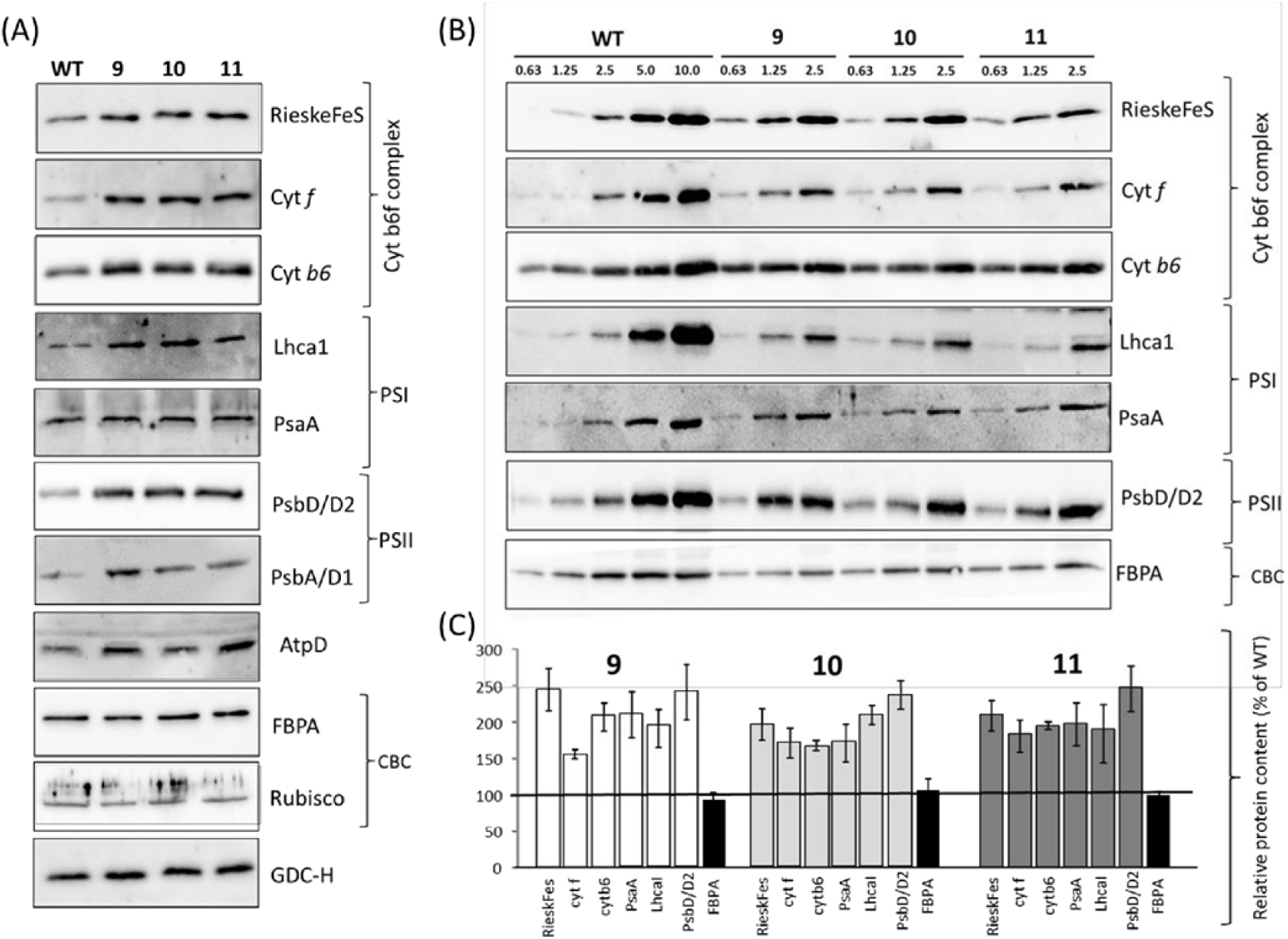
Immunoblot analysis of leaf proteins of wild type and Rieske FeS ox plants. Protein extracts from leaf discs taken from two leaves per plant from three independent lines (9, 10 and 11) and separated on a 12% acrylamide gel, transferred to membranes and probed with relevant antibodies. *cytochrome b6f* complex subunits, Photosystem I (PSI), Photosystem II (PSII), ATP synthase delta subunit (AtpD), Calvin-Benson cycle (CBC) proteins and the photo-respiratory GDC-H protein were probed. (A) Protein (6 μg) was loaded for all antibodies except for FBPA (3 μg) and rubisco (1 μg). (B) Proteins were loaded containing 0.63 to 10 μg of protein. (C) Protein levels were statistically analysed against WT grown plants using a one-sample t-test (* p < 0.05, ** p < 0.01) and presented as relative protein content compared to WT.

### Chlorophyll fluorescence imaging reveals increased photosynthetic efficiency in young Rieske FeS ox seedlings

In order to explore the impact of increased levels of the Rieske FeS protein on photosynthesis the quantum efficiency of PSII (*F*_q_’/*F*_m_’) was analysed using chlorophyll a fluorescence imaging (Baker, 2008; Murchie and Lawson, 2013). A small increase in *F*_q_’/*F*_m_’ was found in the Rieske FeS ox plants at irradiances of 310 μmol m^-2^ s^-1^ and 600 μmol m^-2^s^-1^ (Fig. 2). Leaf area, generated from these images, was significantly larger in all Rieske FeS ox lines compared to WT (Fig. 2C), but no significant difference in leaf thickness was observed between the leaves of Rieske FeS lines 9 and 11 and that of the WT plants (Supplemental Table S1).

**Figure 2.**
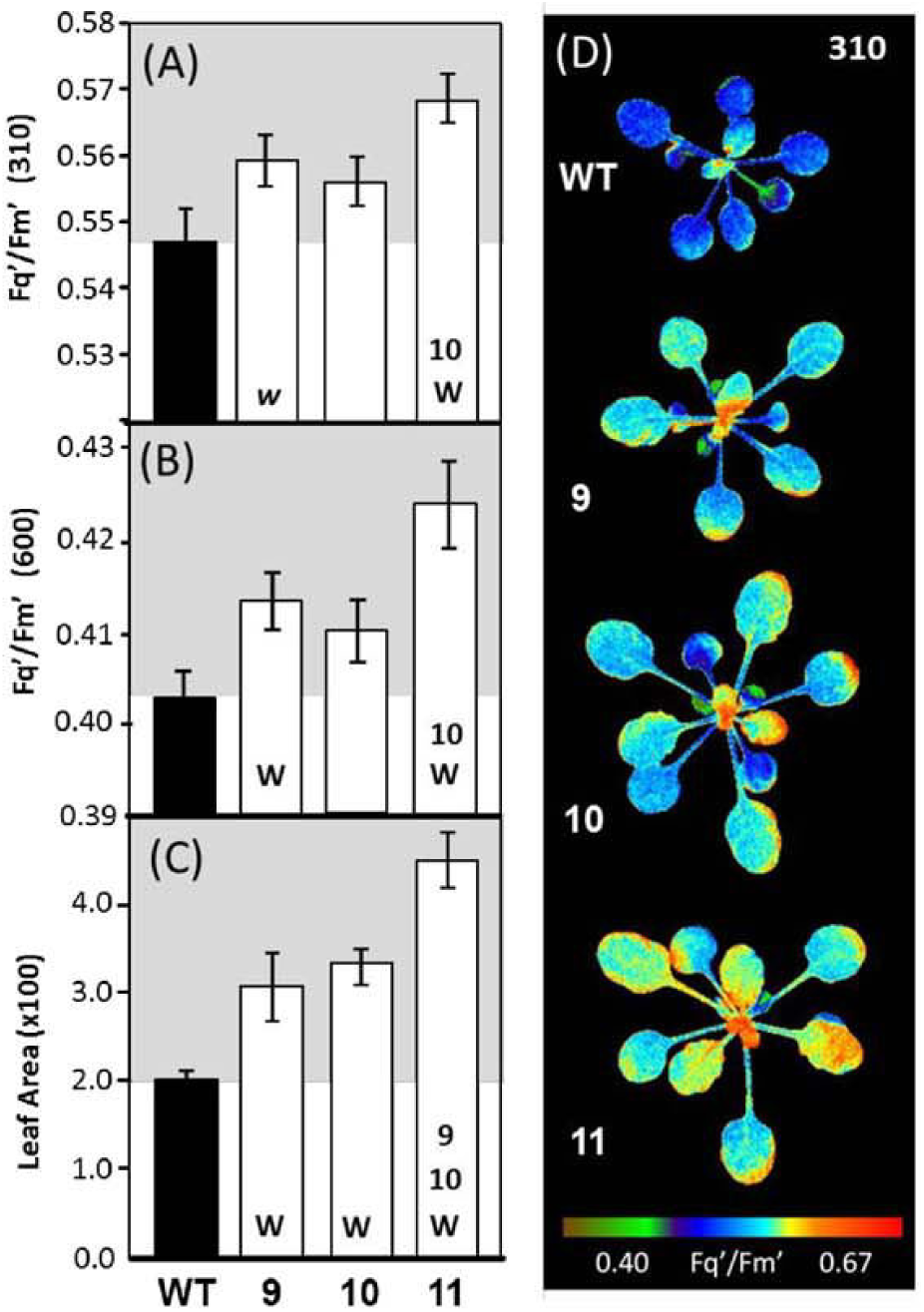
Determination of photosynthetic capacity and leaf area in Rieske FeS ox seedlings using chlorophyll fluorescence imaging. WT and Rieske FeS ox plants were grown in controlled environment conditions with a light intensity of 130 μmol m^-2^ s^-1^, 8 h light/16 h dark cycle and chlorophyll fluorescence imaging used to determine *F*_q_’/*F*_m_’ (maximum PSII operating efficiency) at two light intensities 14 days after planting (DAP) (A, D) Fluorescence *F*_q_’/*F*_m_’ at 310 μmol m^-2^ s^-1^, and (B) *F*_q_’/*F*_m_’ at 600 μmol m^-2^ s^-1^ (C) leaf area at time of analysis. The data was obtained using four to six individual plants from each line compared to WT (five plants). Significant differences (p<0.05) are represented as capital letters indicating differences between line. Bars represent Standard errors. Lower case italic lettering indicates lines are just below significance (p>0.05 - <0.1).

### Photosynthetic CO_2_ assimilation and electron transport rates are increased in the Rieske FeS ox plants

The impact of overexpression of the Rieske FeS protein on the rate of photosynthesis in mature plants was investigated using combined gas exchange and chlorophyll fluorescence analyses. Both the light saturated rate of CO_2_ fixation (*A*_sat_) and the relative light saturated rate of electron transport (rETR), were increased in the Rieske FeS ox lines compared to WT when measured at 2% [O_2_] (Fig. 3A & B; Table 1). Additionally the light saturated rate of CO_2_ assimilation at ambient [CO_2_] was also increased when measured at 21% [O_2_] (Supplemental Fig. S4). No significant difference in leaf absorbance (Abs) between the Rieske FeS ox and WT plants was found (Table S1).

**Figure 3.**
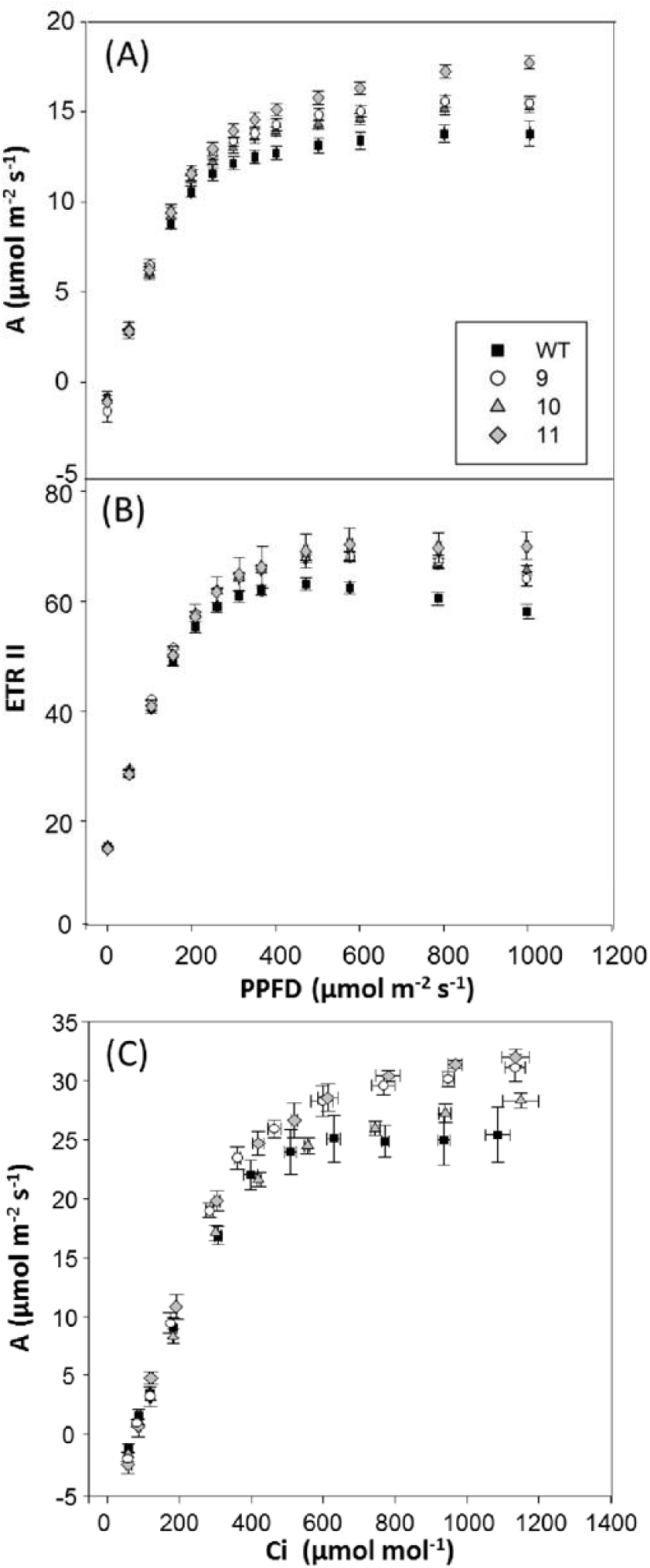
Photosynthetic responses of the Rieske FeS ox plants. (A) Determination of photosynthetic capacity and (B) electron transport rates in transgenic plants at 2% [O_2_]. WT and transgenic plants were grown in controlled environment conditions with a light intensity 130 μmol m^-2^ s^-1^, 8 h light/16 h dark cycle for four weeks, (C) Photosynthetic carbon fixation rate (*A*) was determined as a function of increasing CO_2_ concentrations (*A*/*C*_i_) at saturating-light levels (1000 μmol m^-2^ s^-1^). WT and transgenic plants were grown in controlled environment conditions at a light intensity 280 μmol m^-2^ s^-1^, 12 h light/12 h dark cycle for four weeks. Error bars represent standard errors.

**Table 1.** Photosynthetic parameters of WT and Rieske FeS ox lines determined from light response (*AQ*) curves carried out at 2% [O_2_] (see Fig. 3a and 3b). Statistical differences are shown in bold (* p<0.1; ** p<0.05; *** p<0.01). Standard errors are shown

| Name | **A/Q 2% [O<sub>2</sub>]** |  |  |  |  |
| --- | --- | --- | --- | --- | --- |
| | Light<br>($\mu\text{mol m}^{-2} \text{s}^{-1}$) | $F_q'/F_m'$ | ETR II<br>($\mu\text{mol e}^- \text{m}^{-2} \text{s}^{-1}$) | $q_P$ | $A_{\text{sat}}$<br>($\mu\text{mol m}^{-2} \text{s}^{-1}$) |
| **WT** | 400 | 0.343 ± 0.013 | 62.3 ± 0.96 | 0.58 ± 0.02 |  |
|  | 1000 | 0.124 ± 0.002 | 54.4 ± 0.91 | 0.23 ± 0.01 | 14.7 ± 0.35 |
| **9** | 400 | **0.380 ± 0.004 \*\*** | **66.5 ± 0.46 \*\*** | **0.64 ± 0.00 \*\*** |  |
|  | 1000 | **0.151 ± 0.002 \*\*\*** | **65.9 ± 0.79 \*\*\*** | **0.28 ± 0.00 \*\*** | **16.5 ± 0.13 \*\*** |
| **10** | 400 | 0.369 +/-0.009 | **65.9 ± 0.68 \*\*** | 0.62 ± 0.01 |  |
|  | 1000 | **0.147 ± 0.003 \*\*\*** | **64.3 ± 1.24 \*\*\*** | **0.27 ± 0.00 \*** | **17.7 ± 0.87 \*\*\*** |
| **11** | 400 | 0.370 ± 0.018 | 66.4 ± 3.73 | 0.63 ± 0.02 |  |
|  | 1000 | **0.156 ± 0.006 \*\*\*** | **70.3 ± 2.47 \*\*\*** | **0.29 ± 0.00 \*\*\*** | **18.4 ± 0.44 \*\*\*** |

In plants grown at a light level of 130 μmol m^-2^ s^-1^ no difference in the light- or CO_2_ - saturated rate of CO_2_ assimilation (*A*_max_) was found. In contrast, in a second group of plants grown at 280 μmol m^-2^s^-1^, *A*_max_ was greater in the Rieske FeS ox lines 9 and 11 relative to WT (Fig. 3C; Table 1). Further analysis of the *A*/*C*i curves revealed that *J*_max_ was significantly greater in the Rieske FeS ox plants when compared to WT (Table 1), but no significant difference in *Vc*_max_ (data not shown) was observed.

**Table 2.** Maximum electron transport rate (*J*_max_) and maximum assimilation (*A*_max_) in WT and Rieske FeS ox lines derived from the *A*/*C*_i_ response curves shown in Fig. 3c using the equations published by von Caemmerer and Farquhar (1981). Statistical differences are shown in bold (* p<0.1; ** p<0.05;). Standard errors are shown.

| Name | **A /Ci** |  |
| --- | --- | --- |
| | $J_{\text{max}}$<br>($\mu\text{mol m}^{-2} \text{s}^{-1}$) | $A_{\text{max}}$<br>($\mu\text{mol m}^{-2} \text{s}^{-1}$) |
| **WT** | 181 ± 11.6 | 24.4 ± 2.31 |
| **9** | **210 ± 10.2 \*** | **31.1 ± 1.14 \*\*** |
| **10** | 194 ± 8.9 | 28.3 ± 0.67 |
| **11** | **216 ± 11.9 \*\*** | **31.9 ± 0.68 \*\*** |

### The quantum efficiency of PSI and PSII was increased in the Rieske FeS ox plants

To further explore the influence of increases in the Rieske FeS protein on PSII and PSI photochemistry, dark-light induction responses were determined in WT and Rieske FeS ox (lines 11 & 10) using simultaneous measurements of P700 oxidation state and PSII efficiency. These results showed that the quantum yields of both PSI and PSII were increased in the Rieske FeS ox plants compared to WT and that the fraction of PSII centres that were open (*q*_L_) was also increased, whilst the level of Qa reduction (1-*q*_p_) was lower in leaves of 27 day old plants from line 11. (Fig. 4). NPQ levels were also shown to be lower in the Rieske FeS ox plants together with a reduction in stress induced limitation of NPQ (*q*_N_) when compared to WT plants (Fig. 4). Similar results were obtained for both lines 10 and 11 when plants were analysed later in development (34 DAP) (Supplemental Fig. S5). The increases in the quantum yields of PSI and PSII observed here were accompanied by corresponding increase in electron transport rates (ETRI and ETRII; Supplemental Fig. S6).

**Figure 4.**
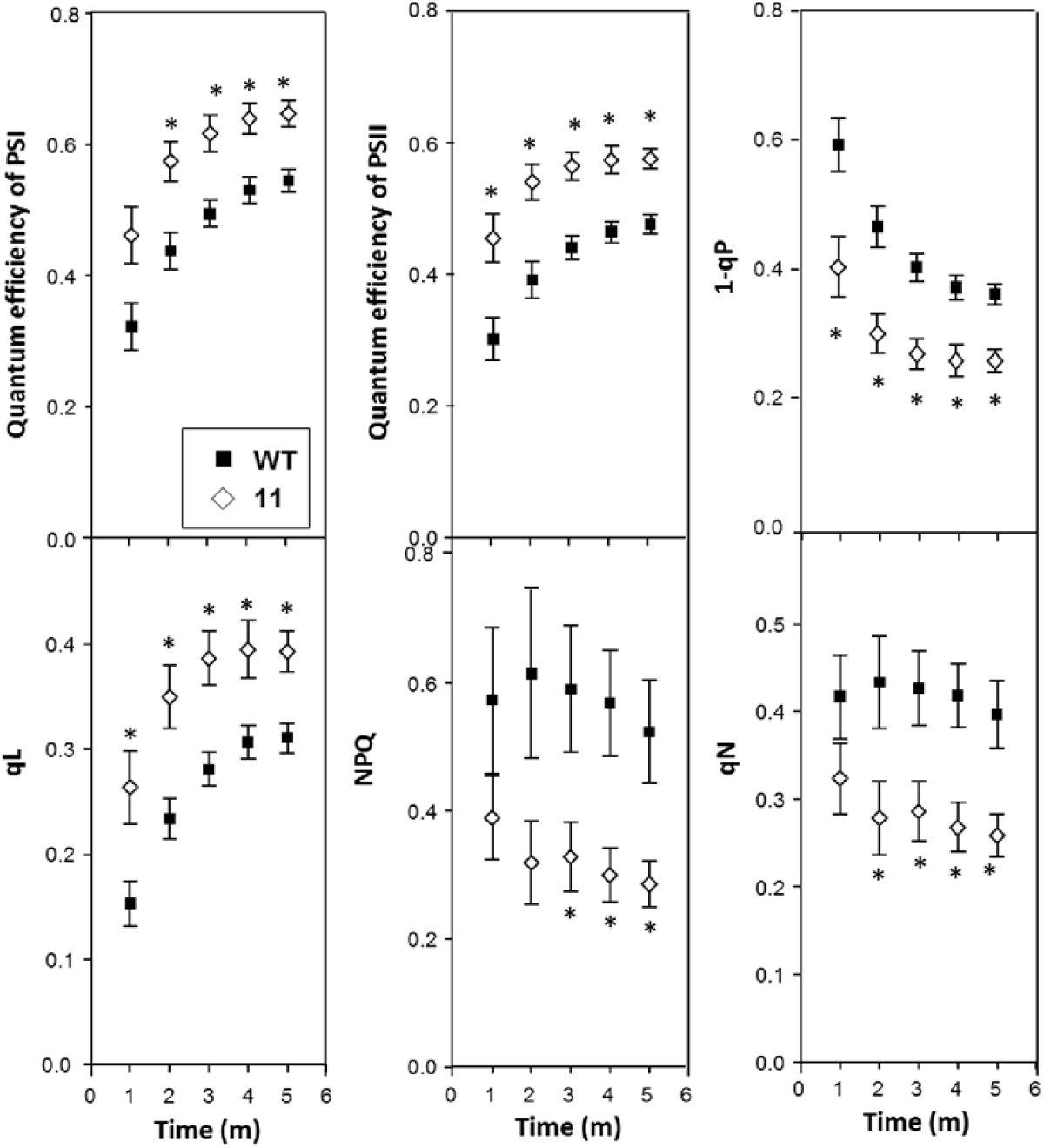
Determination of the efficiency of electron transport in the leaves of young Rieske Fes ox plants. WT and Rieske FeS ox plants were grown in controlled environment conditions with a light intensity of 130 mmol m^-2^ s^-1^, 8 h light/16 h dark cycle and the redox state was determined (27 days after planting) using a Dual-PAM at a light intensity of 220 μmol m^-2^ s^-1^. The data was obtained using four individual plants from Rieske Fes ox line 11 compared to WT (five plants). Significant differences are indicated (*p<0.05). Bars represent standard errors.

### Growth, vegetative biomass and seed yield is increased in the Rieske FeS ox plants

The leaf area of the Rieske FeS ox lines was significantly greater than WT as early as 10 days after planting in soil and by 18 days were 40-114% larger (Fig. 5). Destructive harvest at day 25 showed that this increase in leaf area translated to an increase in shoot biomass of between 29% - 72% determined as dry weight (Fig. 5C). To determine the impact of increased Rieske FeS protein on seed yield and final shoot biomass a second group of plants was grown in the same conditions as described in Fig 6. Interestingly, 38 days after planting (DAP) 40% of the Rieske FeS ox plants had flowered in contrast to 22% in the WT plants (Fig. 6A). Following seed set (52 DAP) both vegetative biomass (Fig. 6B) and seed yield (Fig. 6C) were determined and although a significant increase in biomass was observed in all of the Rieske FeS ox plants a statically significant increase in seed yield was only evident in line 11.

**Figure 5.**
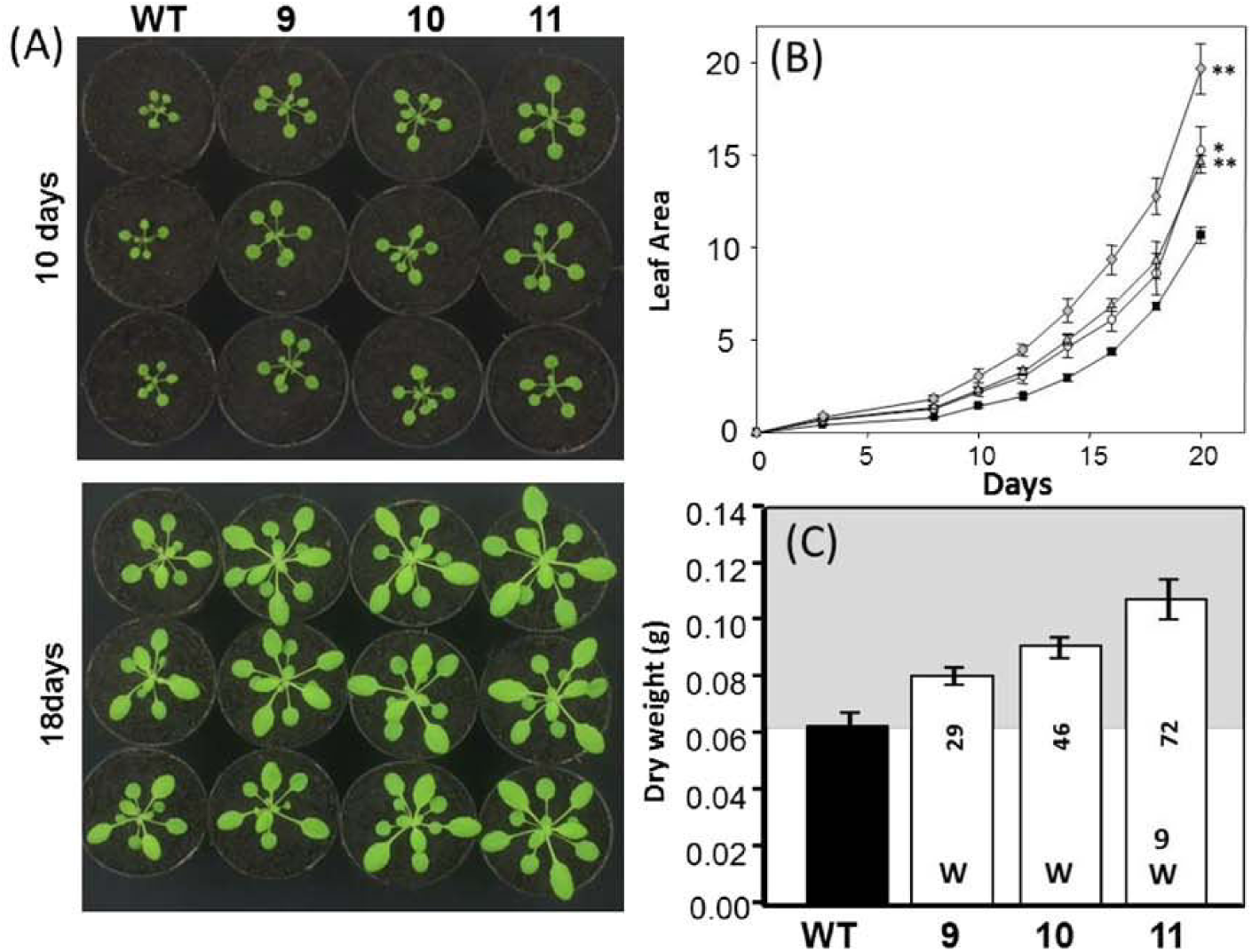
Growth analysis of WT and Rieske FeS ox plants. Plants were grown at 130 μmol m^-2^ s^-1^ light intensity in long days (8 h/16 h days). (A) appearance of plants at 10 and 18 days after planting (DAP). (B) Leaf area determined 20 DAP. (C) Final biomass at 25 DAP. Results are representative of six to nine plants from each line. Increase over WT (%) is indicated as numbers on histogram. Significant differences (p<0.05) are represented by capital letters. Significant differences * (p<0.01), ** (p<0.001), are indicated. Bars represent standard errors.

**Figure 6.**
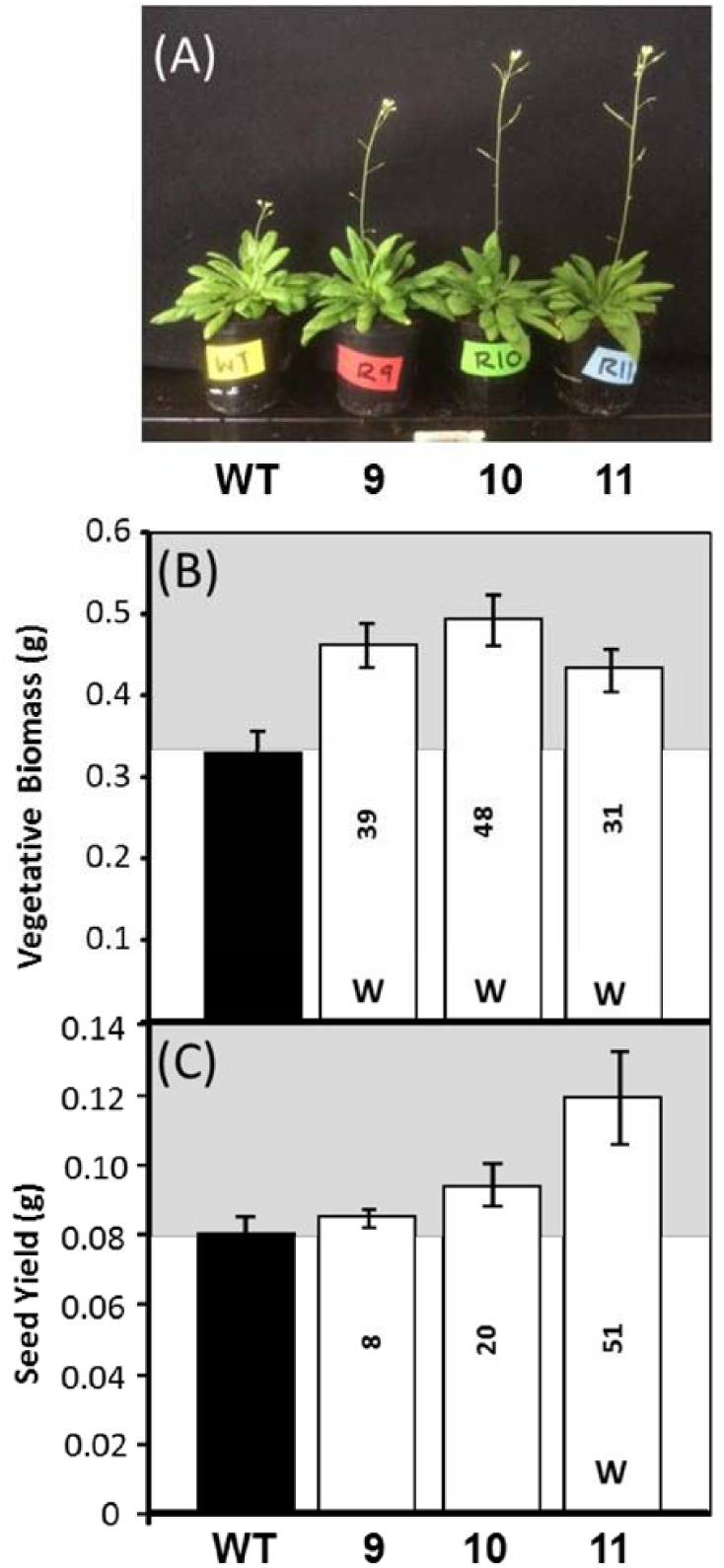
Seed yield and vegetative biomass of WT and Reiske FeS ox plants. Plants were grown at 130 μmol m^-2^ s^-1^ light intensity in long days (8 h/16 h days). (A) Appearance of plants at 38 DAP. (B) final biomass and (C) seed yield at harvest (52 DAP). Increase over wild type (%) is indicated by numbers on histogram. Results are representative of six to nine plants from each line. Significant differences (p<0.05) are represented by capital letters. Bars represent standard errors.

### The pigment content was altered in the Rieske FeS ox plants

The pigment composition of the leaves of the Rieske FeS ox and WT plants was determined. An increase in the levels of chlorophyll a and b (14-29%) was observed in the Rieske FeS ox compared to WT plants (Fig. 7). These increases were accompanied by an increase in the carotenoids, neoxanthin (+38%), violaxanthin (+59%), lutein (+75%) and β-carotene (+169%). No detectable change in the level of zeaxanthin was evident in the Rieske FeS ox plants.

**Figure 7.**
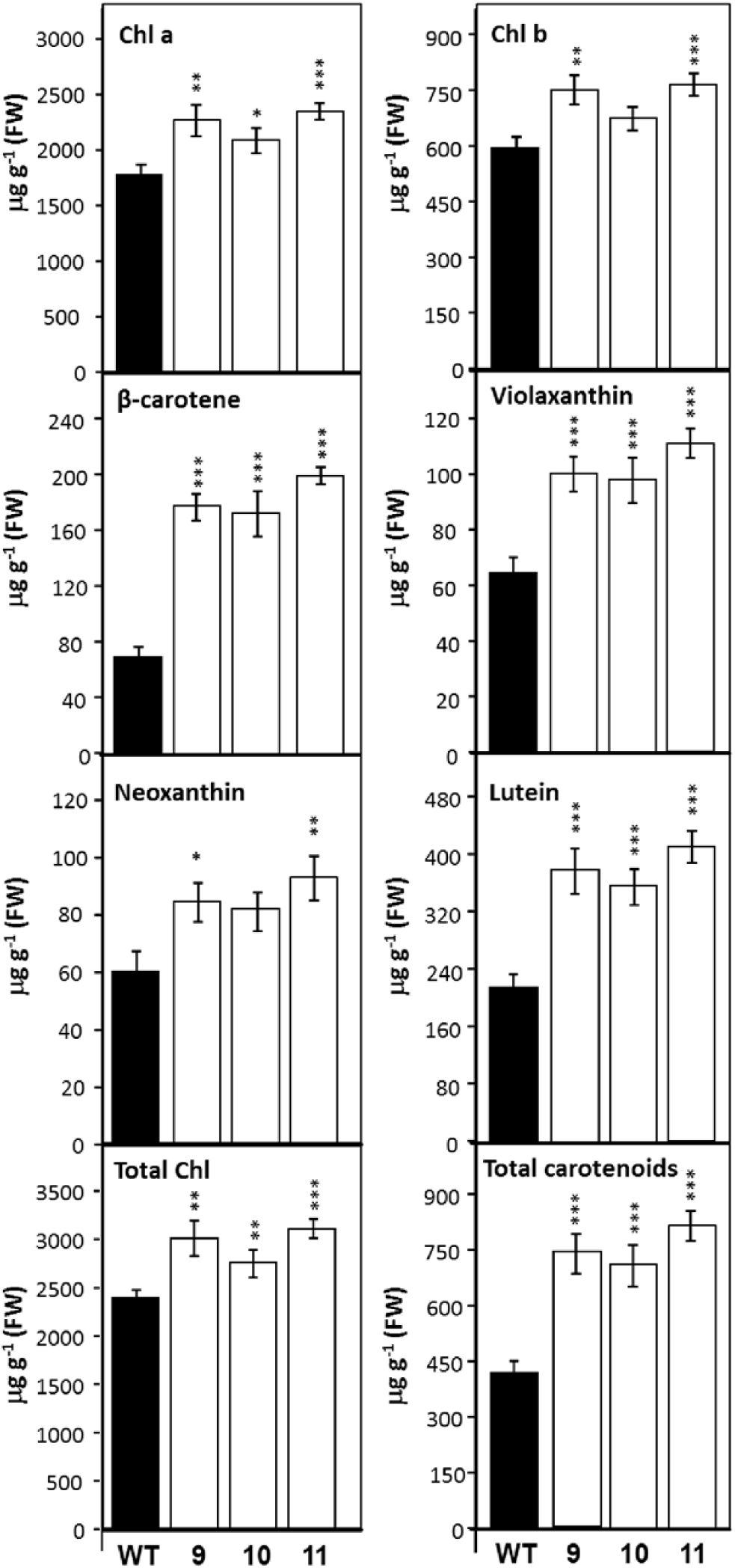
Pigment content in WT and Rieske FeS ox plants. Plants were grown at 130 μmol m^-2^ s^-1^ light intensity in short days (8h/16 h days). Two leaf discs, collected from two different leaves, were immersed in DMF at 4°C for 48 hours and separated by UPLC. Results are represented as μg/g^-1^ fresh weight. Statistical differences are shown in bold (* p<0.1; ** p<0.05; *** p<0.001). Bars represent standard errors.

### Discussion

In recent years increasing the rate of photosynthetic carbon assimilation has been identified as a target for improvement to increase yield. Evidence to support this has come from the theory and modelling of the photosynthetic process, growth of plants in elevated CO_2_ and also from transgenic manipulation (Zhu et al., 2010). It was shown previously in antisense studies that reducing the levels of the Rieske FeS protein resulted in a reduction in levels of the cyt *b*_*6*_*f* complex, a decrease in photosynthetic electron transport and in rice a decrease in both biomass and seed yield was observed (Price et al., 1998; Yamori et al., 2016). These findings identified the cyt *b*_*6*_*f* complex as a limiting step in electron transport and would suggest that over expression of the Rieske FeS protein may be a feasible route to increase photosynthesis and yield. In this study we show that overexpression of the Rieske FeS protein in Arabidopsis results in an increase in photosynthesis, vegetative biomass and seed yield.

### Increased levels of the Rieske FeS protein increased photosynthetic electron transport, CO_2_ assimilation and biomass

Using chlorophyll fluorescence imaging we have shown that that overexpression of the Rieske FeS protein resulted in an increase in photosynthesis and growth which is evident from the early stages of seedling development. These observed increases in *F*_q_’/*F*_m_’ represent an early indication that the potential quantum yield of PSII photochemistry had been elevated in Rieske ox lines (Genty et al., 1989; Genty et al., 1992; Baker et al., 2007). This early stimulation is maintained into maturity and an increase in the light saturated rate of CO_2_ assimilation and electron transport rates was evident in the Rieske FeS ox plants. Our data also showed that quantum yields of both PSII and PSI were increased and that the fraction of PSII centres available for photochemistry was increased indicated by an increase in (*q*_L_) and a lower 1-*q*_p_ (Baker et al., 2007; Baker and Oxborough, 2004; Kramer et al., 2004). These results are consistent with what would be predicted from results obtained from the Rieske FeS antisense studies where ETR was reduced (Price. et al., 1998; Ruuska et al., 2000; Yamori et al,. 2011a). However, the impact of overexpression of the Rieske FeS protein was clearly not restricted to increasing the activity of the cyt *b*_6_/*f* complex but resulted in an increase in electron flow through the entire electron transport chain.

Substantial and significant increases in the growth of the rosette area were observed in the Rieske FeS ox plants in the early vegetative phase which resulted in an approximately 30-70% increase in biomass yield in the different lines. Importantly seed yields in line 11, which showed the biggest increases in shoot biomass were also shown to be increased relative to WT.

### The Rieske FeS ox plants have increased levels of proteins in the cyt *b*_*6*_*f*, PSI and PSII complexes

In keeping with our analysis of the electron transport processes in the Rieske FeS ox plants, an increase in the levels of the cyt *b*_6_ and cyt *f*, core proteins of the cyt *b*_*6*_*f* complex was evident. Furthermore an increase in the levels of proteins in both PSII and PS1 and the δ subunit of the ATPase complex was also observed. This result was unexpected given that no changes in components of PSII or PSI were observed in the Rieske FeS antisense plants. Interestingly, a recent study reported increases in cyt *b*_6_*f* proteins levels in Arabidopsis plants grown under square wave light, compared to plants grown under fluctuating light, and these were matched by increased levels of PSII, PSI and the δ subunit of the ATPase proteins, which agrees with our study (Vialet-Chabrand et al., 2017). Furthermore, the *hcf* mutant, in which the biogenesis of the cyt *b*_*6*_*f* was inhibited, had a reduced accumulation of components of both PSI and PSII, although these complexes remained fully functional, as inferred from spectroscopic analyses, and no mechanism controlling these changes has been identified (Lennartz et al., *2001*). Despite considerable efforts to gain insight into the mechanisms underlying the regulation of synthesis and assembly of components of the thylakoid membrane, the factors determining accumulation of these complexes are still poorly understood but our results provide evidence that the Rieske FeS protein levels may play a role in this regulation (Schöttler, 2015).

### Over-expression of Rieske FeS significantly modifies pigment content of leaves

In parallel with the increase in components of the thylakoid membranes, plants with increased Rieske FeS protein were found to have an increase in levels of both chlorophyll a and b and a small increase in the chlorophyll a to b ratio from 2.96 to 3.12. The increase in chlorophyll a and b suggest a greater investment in both light capture and PSII reactions centres and would fit with the increase in photosynthetic electron transport capacity in the Rieske FeS ox plants. In previous work, plants with reduced levels of the Rieske FeS protein had a lower chlorophyll *a*/*b* ratio (Hurry et al., 1996; Price et al., 1998) which is the opposite to what was observed in the Rieske FeS ox plants. In addition to increases in chlorophyll, significant increases in the carotenoid pigments were also seen with β-carotene, violaxanthin (+59%), lutein (+75%) and neoxanthin (+37%). β-carotene is a component of both the RC and light harvesting complex (Kamiya and Shen, 2003; Ferreira et al., 2004; Loll et al., 2005; Litvin et al., 2008; Janik et al., 2016) and the increase in these pigments observed in the Rieske FeS ox plants is in agreement with investment in both light harvesting and increasing RC efficiency.

Lutein, neoxanthin and violaxanthin are the main xanthophyll pigment constituents of the largest light-harvesting pigment-protein complex of photosystem II (LHCII) (Thayer and Björkman, 1992; Ruban et al., 1994; Ruban et al., 1999; Ruban et al., 2012; Janik et al., 2016). Acidification of the thylakoid lumen as a result of electron transport (and driven in particular by the activities of the cytochrome *b*_*6*_*f* complex) is accompanied by the de-epoxidation of violaxanthin and an accumulation of zeaxanthin (Björkman and Demmig-Adams, 1994; Müller et al., 2001; Ruban et al., 2012), as well as protonation of carboxylic acid residues of the PsbS protein associated with PSII antennae (Li et al., 2000, 2004). Protonation of PsbS and binding of zeaxanthin increase NPQ and the thermal dissipation of excitation energy (Baker, 2008; Jahns and Holzwarth, 2012). Increases in electron transport observed in Rieske FeS ox lines might be expected to result in the acidification of the thylakoid lumen and an increase in NPQ. However, we found that the increase in the level of the Rieske FeS protein led to a small but significantly lower steady state levels of NPQ and an increase in the rate of relaxation of NPQ following illumination. The absence of an increase in NPQ in the presence of significant increases in electron transport rates suggest that the Rieske FeS ox plants also have increased rates of ATP synthesis. Although we provide no direct support for this we did observe and increase in the level of the ATP synthase delta subunit protein in the Rieske FeS ox plants. Support for this comes from the earlier work on the Rieske FeS antisense plants showing that the levels of ATP synthase were reduced and that a low transthylakoid pH gradient was evident (Price et al., 1995, 1998 Ruuska et al., 2000).

### Conclusion

A number of studies have shown that increasing photosynthesis through the manipulation of CO_2_ assimilation can improve growth (Miyagawa et al., 2001; Lefebvre et al., 2005; Rosenthal et al., 2011; Uematsu et al., 2012; Simkin et al., 2015, 2017), this work together with a study in which cytochrome C_6_ from the red alga *Porphyra*, was expressed in Arabidopsis (Chida *et al*., 2007) provide direct evidence that there is also an opportunity to improve the efficiency of the electron transfer chain. Our results demonstrate that overexpression of the Rieske FeS protein can increase electron transport, photosynthesis and yield and provides another potential avenue to improve crop productivity and meet the food requirements for future population growth. Furthermore, overexpression of the Rieske FeS protein may offer tool to investigate fundamental questions on factors controlling the biogenesis of the photosynthetic complexes in the electron transport chain.

## ACKNOWLEDGMENTS

We thank James E. Fox and Philip A. Davey for help with pigment analysis and Elena A. Pelech for help with Dual-PAM measurements. A.J.S was supported by BBSRC (Grant: BB/J004138/1 awarded to C.A.R and T.L): A.J.S generated transgenic plants and performed molecular and biochemical experiments and carried out plant phenotypic and growth analysis. A.J.S and L.M performed gas exchange measurement on Arabidopsis. A.J.S and L.M carried out data analysis on their respective contributions. C.R and T.L designed and supervised the research. C.R, A.J.S wrote the manuscript with input from TL.

## Supplementary Information

**Figure S1.**
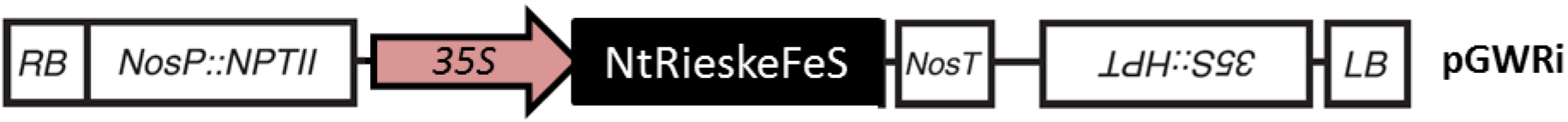
Schematic representation of the RieskeFeS over-expression vector pGWRi used to transform Arabidopsis (Col-0). cDNA are under transcriptional control of the 35s tobacco mosaic virus promoter (*35S*), which directs constitutive high-level transcription of the transgene, and followed by the *nos* 3' terminator (*NosT*). Following floral dipping, transgenic Arabidopsis plants were selected on both kanamycin (*NPTII*) and hygromycin (*HPT*) containing medium (Nakagawa *et al.* 2007). *RB*; T-DNA right border, *LB*; T-DNA left border, *NosP*; nopaline synthase promoter,.

**Figure S2.**
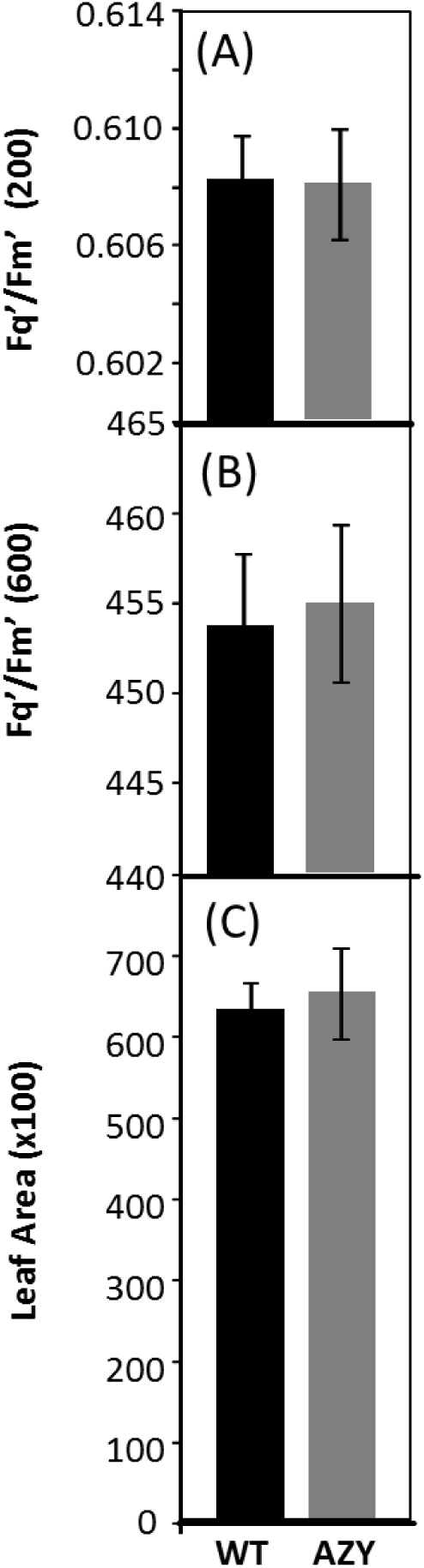
Determination of photosynthetic capacity and leaf area of WT versus azygous (AZY) segregating controls using chlorophyll fluorescence imaging. WT and AZY plants were grown in controlled environment conditions with a light intensity of 130 μmol m^-2^ s^-1^, 8 h light/16 h dark cycle for 14 days and chlorophyll fluorescence imaging used to determine *F*_q_’/*F*_m_’ (maximum PSII operating efficiency) at a light intensity of (A) 200 μmol m^-2^ s^-1^. (B) 600 μmol m^-2^ s^-1^ (The data was obtained using 4 individual plants from each line). (C) leaf area at time of analysis.

**Figure S3.**
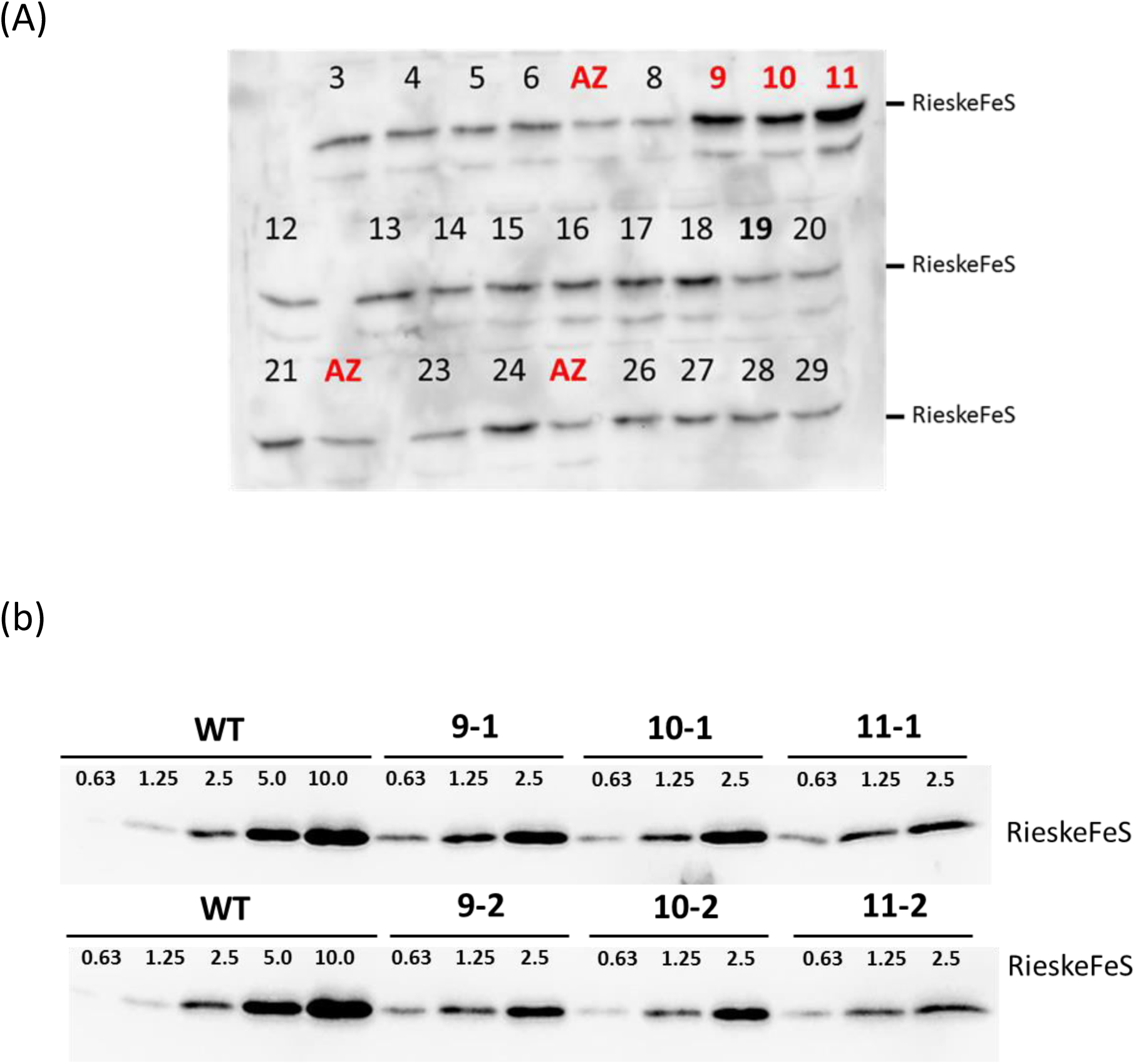
Immunoblot analysis of WT and Rieske FeS ox proteins. (A) Leaf protein extract from leaves from 27 independent T1 transformants. Protein levels were compared to PCR negative lines (AZ). Lines 9, 10 and 11 were selected for further analysis. (B) Protein gradients were loaded for two independent plants per line containing 0.63 to 10μg of protein.

**Figure S4.**
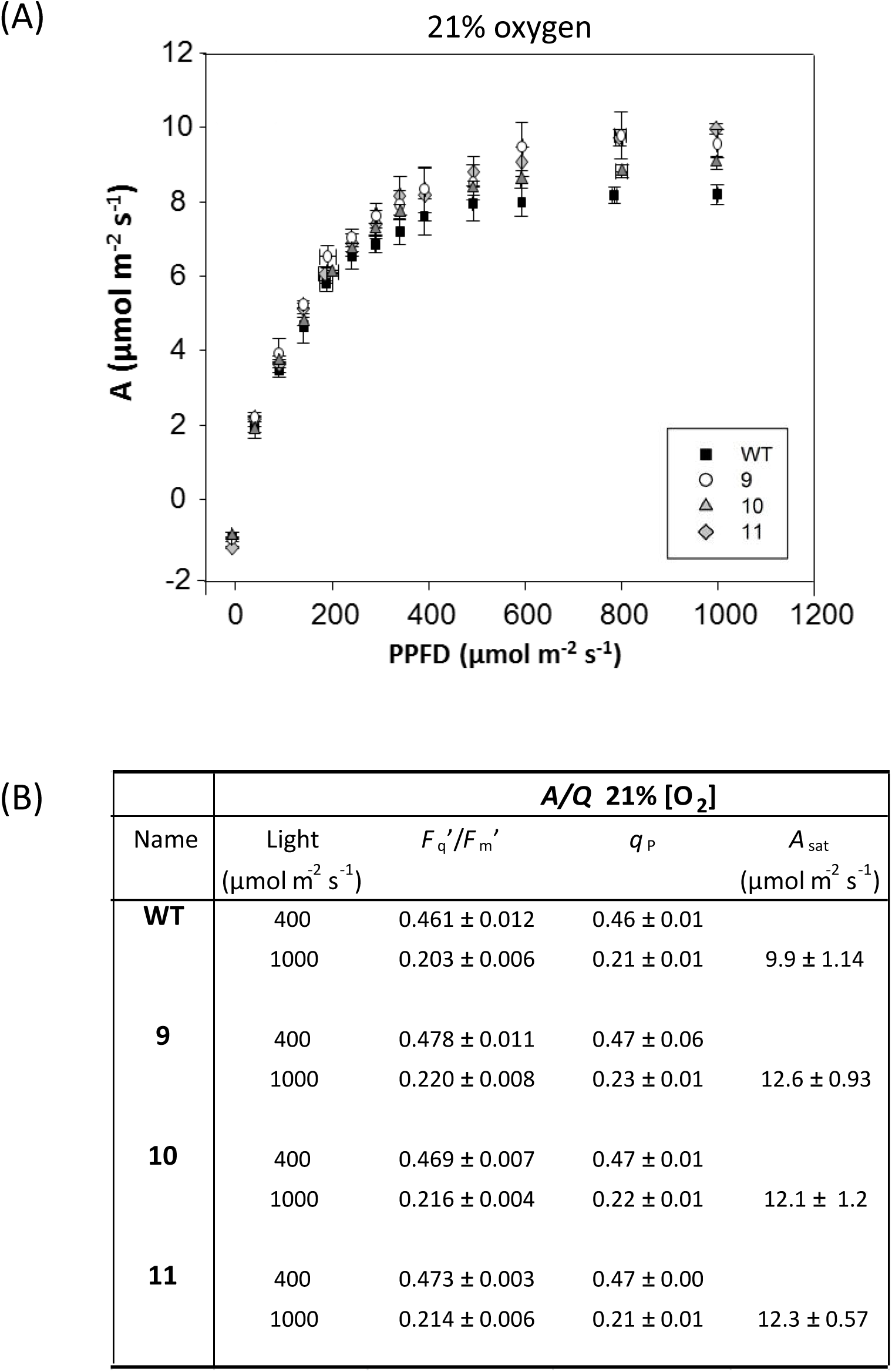
Light Response curves of the Rieske FeS plants at 21% [O_2_]. (A) Light Response Curves for each Rieske FeS lines compared to wild type (WT). The data was obtained using four individual Rieske FeS ox and WT plants. Standard errors are shown. (B) the photosynthetic parameters of WT and Rieske FeS ox lines determined from light response (*AQ*) curves.

**Figure S5.**
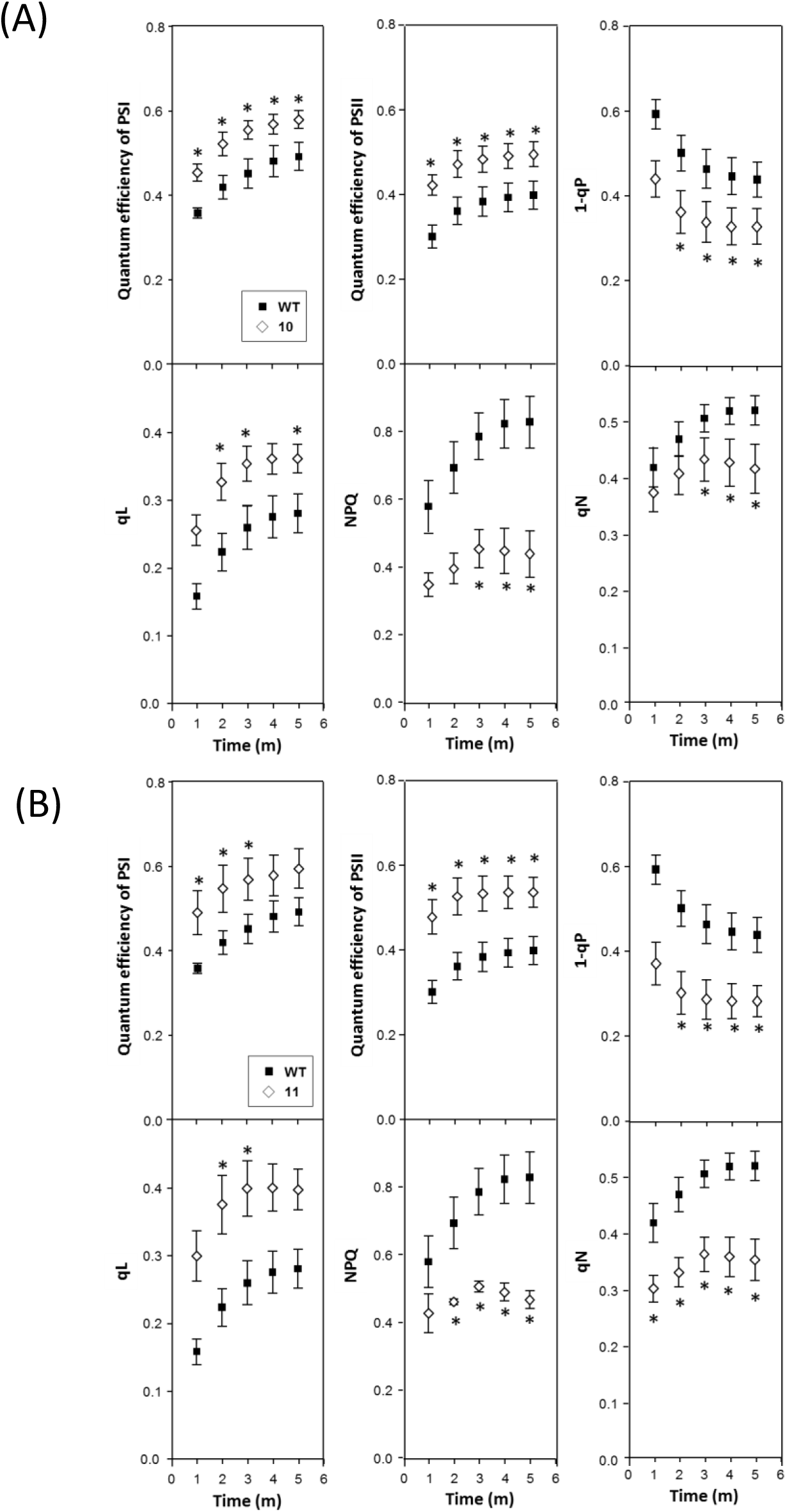
Determination of the efficiency of electron transport in the leaves of mature Rieske Fes ox plants (lines 10 & 11, 34 days after planting). WT and Rieske FeS ox plants were grown in controlled environment conditions with a light intensity of 130 μmol m^-2^ s^-1^, 8 h light/16 h dark cycle and the redox state was determined using a Dual-PAM at a light intensity of 220 μmol m^-2^ s^-1^. The data was obtained using four individual plants from Rieske Fes ox line (A) 10 and (B) 11 compared to WT (five plants). Significant differences are indicated (*p<0.05). Bars represent Standard errors.

**Figure S6.**
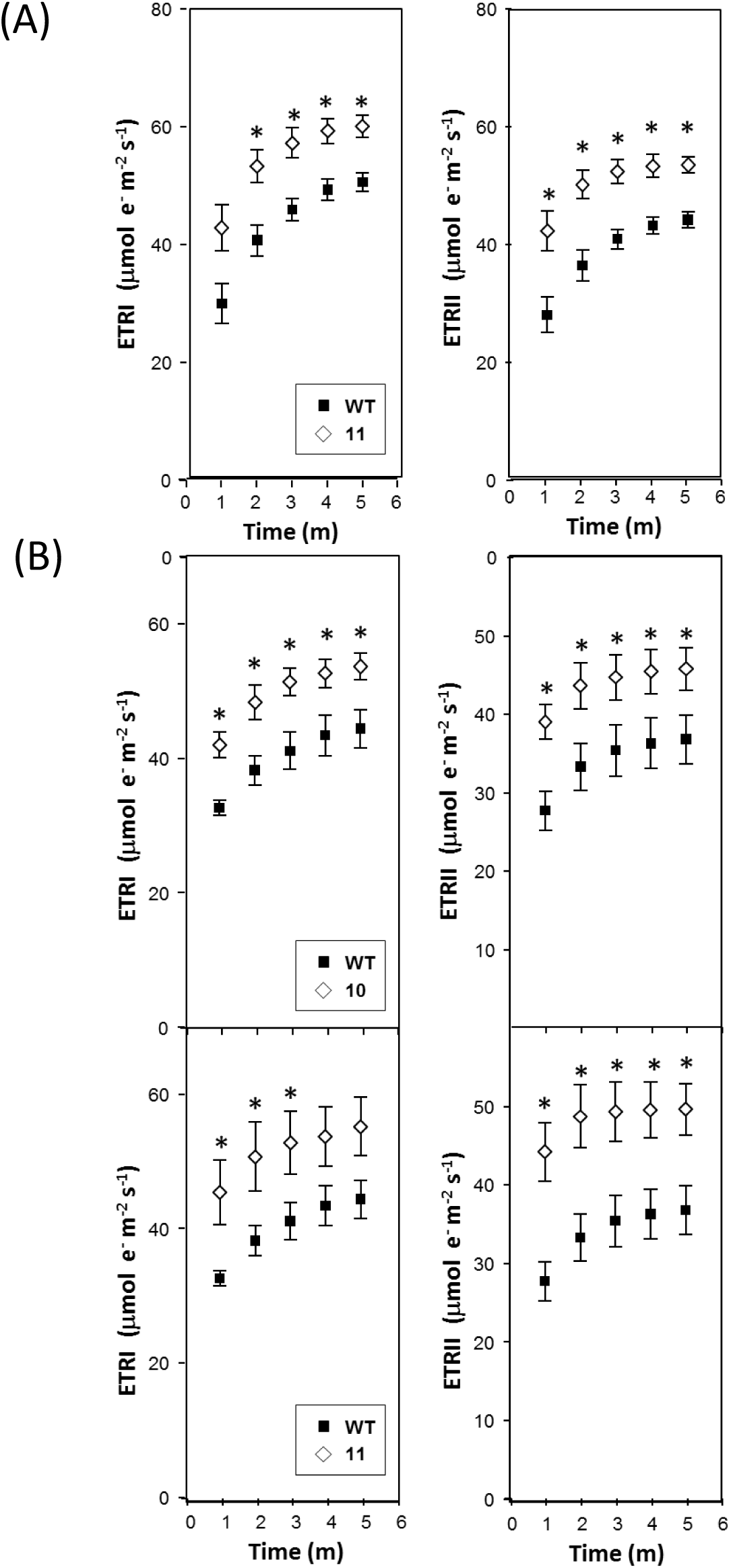
Determination of the efficiency of electron transport in the leaves of (A) young (line 11, (27 days after planting)) and (B) mature Rieske Fes ox plants (lines 10 & 11, (34 days after planting respectively)). WT and Rieske FeS ox plants were grown in controlled environment conditions with a light intensity of 130 μmol m^-2^ s^-1^, 8 h light/16 h dark cycle and the redox state was determined using a Dual-PAM at a light intensity of 220 μmol m^-2^ s^-1^. The data was obtained using four individual plants from Rieske Fes ox line Significant differences are indicated (*p<0.05). Bars represent Standard errors.

**Table S1.** Physiological parameters of Rieske FeS ox plants compared to WT. The data was obtained using four individual Rieske FeS ox and WT plants. Standard errors are shown. WT and Rieske FeS ox plants were grown in controlled environment conditions with a light intensity of 130 μmol m^-2^ s^-1^, 8 h light/16 h dark cycle

| Plant Type | Abs | $F_v/F_m$ | leaf thickness<br>($\mu\text{m}$) |
| --- | --- | --- | --- |
| WT | $0.878 \pm 0.004$ | $0.789 \pm 0.006$ | $217 \pm 6.2$ |
| 9 | $0.873 \pm 0.015$ | $0.796 \pm 0.009$ | $223 \pm 7.5$ |
| 10 | $0.876 \pm 0.006$ | $0.794 \pm 0.007$ | ----- |
| 11 | $0.881 \pm 0.003$ | $0.793 \pm 0.008$ | $223 \pm 2.5$ |

## Notes

This work was supported by the Biotechnology and Biological Sciences Research Council (Grant: BB/J004138/1 awarded to C.A.R and T.L)

